# Development of human skin equivalents with inducible ceramide depletion for in vitro modelling of lipid deficiency

**DOI:** 10.1101/2025.01.20.633956

**Authors:** Durotimi O. Dina, Miriam Maiellaro, Emanuela Camera, John T. Connelly

## Abstract

The lipid composition of the epidermis plays a critical role in the skin’s barrier function, and defects in lipid synthesis or assembly can cause a spectrum of skin diseases, ranging from dry skin to severe ichthyoses. The aim of this study was to develop an in vitro model of human skin with tuneable lipid deficiency. Human N/TERT keratinocytes were engineered to express doxycycline-inducible short hairpin RNAs targeting ceramide synthase 3 (CerS3), which is essential for synthesis of ultra long chain ceramides and skin barrier function. We show that 3D human skin equivalents (HSEs) with induced knockdown of CerS3 display normal stratification and terminal differentiation but have reduced Nile Red staining for polar lipids. Further analysis of the lipidome by mass spectrometry confirmed a significant reduction in specific classes of ceramides and ceramide chain length in the CerS3 depleted HSEs. We also show that CerS3 knockdown is reversible upon removal of doxycycline and can be used to study recovery and repair of epidermal lipids. Together, these findings provide an overall strategy for genetically tuning the lipid composition within human skin models and establish a new in vitro model of ceramide deficiency.

## Introduction

The skin is a critical first line of defence against pathogens, physical/chemical damage, and ultraviolet radiation, and the uppermost layer of the epidermis, the stratum corneum, forms a selectively permeable barrier that is crucial for the skin’s protective function^1^. The stratum corneum relies on an intricate network of proteins and lipids, including the lipid lamellar bilayer (LLB) and cornified lipid envelope (CLE). The LLB confers the water-impermeable properties of the epidermis and the CLE acts as a sealant for the cohesion of the corneocytes and a scaffold for the LLB^1–5^. These lipid-based structures consist of ceramides, cholesterol and fatty acids, with ceramides making up the greatest fraction by mass^6^.

The specific class of ω-esterified ultra-long chain (ULC) ceramides, also known as the acylceramides (acylCers), are key for the molecular organisation and subsequent function of the LLB and CLE, and the enzyme ceramide synthase 3 (CerS3) is uniquely required for acylCer synthesis^7,8^. To date, five species of ω-esterified hydroxy ceramides have been identified: Cer[EOH], Cer[EOP], Cer[EOS], Cer[EODS] and Cer[EOSD]. Their unique chemical structure, namely their esterification to linoleic acid, is responsible for the formation and organisation of the LLB. Both Cer[EOS] and Cer[EOH] play important roles in the Long Periodicity Phase (LPP) lamellar organisation of the LLB, and it has been shown that in the absence of Cer[EOS] almost no LPP is formed^9–11^. Overall deficiencies in ceramides can compromise the barrier function of the skin and contribute to numerous skin conditions, ranging from dry skin to severe ichthyoses^12–15^, and LLP malformation directly correlates with an inadequate epidermal barrier and a dry skin phenotype^13,16,17^.

CerS3 deficiency in mice leads to a loss of ULC ceramides and subsequent fatality shortly after birth^20^. Mutations in the *CERS3* gene in humans affect barrier function and can cause specific forms of autosomal recessive congenital ichthyoses (ARCI) ^18,19^. For example, the codon exchange mutation (p.Trp15Arg) results in moderate lamellar ichthyosis with a reduction in long chain acylCers^21^, and similar findings have been reported for other mutations leading to Cers3 deficiency^22^. Thus, Cers3 and acylCers appear to be crucial for the lipid barrier function of the skin.

Human skin equivalents (HSEs) that replicate the three-dimensional (3D) structure and function of the skin in vitro are powerful tools for investigating human-specific disease mechanisms and in a few cases have been applied to lipid associated diseases. For example, HSEs constructed using patient-derived keratinocytes have been used to replicate disease phenotypes of CerS3 deficiency in ARCI^21^. HSEs have also been used to model Harlequin Ichthyosis using keratinocytes with engineered knock out of the lipid transporter ATP binding cassette A12 (ABCA12)^23^. Importantly, HSEs are also compatible with lipidomic analysis via mass spectrometry^24^ and can be used to replicate key differences in the lipid composition of the epidermis in diseases, such as atopic dermatitis^25^. However, despite these advances and the importance of lipids in the skin barrier, there are limited experimental models that support precise modulation of lipid synthesis.

As lipid deficiencies can cause a spectrum of conditions with varying degrees of severity, the ability to precisely tune the amount and composition of lipids within an HSE model would be advantageous for in vitro modelling of human lipid disorders and testing drugs and skincare products. In this study, we developed an inducible HSE model of ceramide deficiency using engineered keratinocytes with doxycycline-inducible knockdown (KD) of CerS3. We focused on CerS3 as a proof of concept as it is a critical regulator of ceramide synthesis in the epidermis and is solely responsible for the synthesis of the acylCer class of lipids. We show that reduction of CerS3 expression in keratinocytes results in a global depletion of polar lipids within the epidermis of the HSE, as well as a lower abundance of specific classes of ceramides and a reduction in ceramide chain length. Moreover, we demonstrate the reversibility of inducible CerS3 KD and ability to study the kinetics of lipid recovery within the HSEs following acute ceramide depletion. Together, these findings establish a robust and tuneable methodology for modelling lipid deficiencies in human skin.

## Results

### Establishment of HSEs with inducible CerS3 knockdown

To develop an in vitro human skin model with tuneable control of epidermal lipids we constructed HSEs using keratinocytes with inducible depletion of ceramides, which are essential for maintaining skin moisture and barrier integrity ^2,26,27^. N/TERT keratinocytes were engineered to stably express short-hairpin RNAs (shRNAs) targeting *CERS3* under the control of a doxycycline-inducible promoter^28–30^, alongside constitutive expression of red fluorescence protein (RFP) (Figure 1A). HSEs were constructed by embedding primary dermal fibroblasts in a hydrogel derived from decellularised extracellular matrix of porcine skin (dECM)^31^, and engineered keratinocytes expressing *CERS3* targeting or non-targeting control (NTC) shRNAs were seeded on the surface. HSEs were cultured at the air-liquid interface for 14 days to induce epidermal differentiation and stratification (Figure 1B), and through the addition of doxycycline (Dox) to the culture medium we aimed to induce CerS3 depletion and disruption of the lipid barrier (Figure 1C).

**Figure 1:**
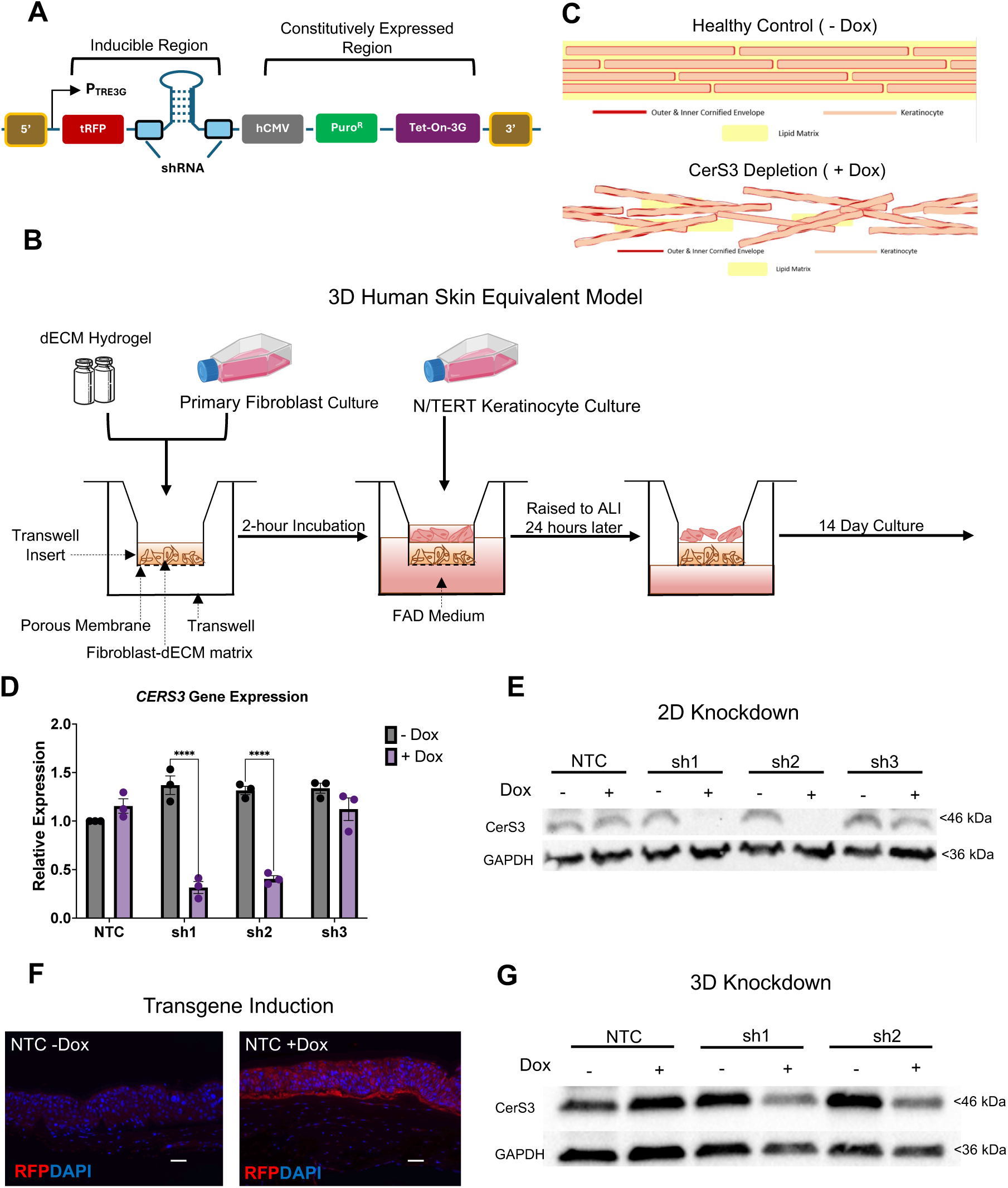
Development of HSEs with inducible CerS3 knockdown. (A) Schematic of the shRNA lentiviral construct (SMARTvector, Horizon Discovery). The constitutively expressed region under the control of the human cytomegalovirus promoter included elements for expression of the Tet-On-3D transactivator and puromycin resistance, while the inducible region contained the shRNA element and tRFP reporter under the control of the TRE3G promoter that is activated by doxycycline (Dox). (B) Schematic overview of the process for constructing 3D HSEs at the air-liquid interface (ALI) of Transwell inserts. (C) Schematic of expected disruption of epidermal lipids following addition of Dox and knockdown of CerS3. (D) Analysis of *CERS3* knockdown in 2D culture by RT-qPCR. Data represent fold-change in expression (2-ΔΔCt method) relative to GAPDH and untreated NTC cells for the shRNA-transduced N/TERT lines without or with Dox treatment (0.1 μg/ml) for 7 days. Data are presented as mean +/− SEM. ****P<0.0001 with 2Way ANOVA and Šidák’s multiple comparison test. N=3 experiments. (E) Representative Western blot for CerS3 and GAPDH expression for all cell lines in 2D culture, without or with Dox treatment for 7 days. (F) Immunofluorescence staining for the inducible tRFP reporter gene (red) in HSEs constructed with NTC keratinocytes without or with Dox treatment for 14 days. Scale bar = 50 μm. (G) Representative Western blot for CerS3 and GAPDH in HSEs constructed with NTC, sh1, and sh2 cell lines and cultured without or with Dox for 14 days.

Analysis of RFP expression in 2D culture first confirmed inducible transgene activation in transduced cells. Maximal RFP expression was reached within 3 days of treatment with a minimum of 0.1 μg/ml of Dox, which was then used for all subsequent experiments (Supplementary Figure S1A). In all four cell lines, over 90% of cells were RFP positive following Dox treatment, while RFP expression was negligible in the absence of Dox (Supplementary Figure S1B). RT-qPCR analysis indicated significant *CERS3* downregulation with Dox exposure in the sh1 and sh2 lines, but not the NTC and sh3 lines (Figure 1D), and a similar KD was observed at the protein level by Western blotting (Figure 1E). As the sh3 line showed no effective KD of CerS3, it was excluded from further analysis.

We next assessed CerS3 KD in 3D HSE models. We confirmed RFP transgene activation through the full thickness of the epidermal layer in the Dox treated cultures (Figure 1F). In accordance with the 2D data, a significant reduction in the CerS3 protein was observed for sh1 and sh2 HSEs treated with Dox compared to untreated controls, but not for the NTC HSEs (Figure 1G, 6B-C). With these results we confirmed KD efficacy and the ability to inducibly deplete CerS3 within the 3D HSE model using two different shRNA constructs, sh1 and sh2.

### CerS3 depletion reduces polar lipid content in HSEs independently of epidermal differentiation and stratification

To determine the effects of CerS3 depletion in the HSE constructs, we first examined overall changes in epidermal structure and terminal differentiation. Hemotoxylin and eosin (H&E) staining indicated no apparent differences in stratification or cornification in the Dox treated sh1 and sh2 models compared to their untreated controls or NTC models (Figure 2A). Likewise, there were no differences in epidermal thickness between any of the conditions (Supplementary Figure S2). Immunofluorescence staining for transglutaminase-1 and loricrin also revealed no discernible differences in expression of late terminal differentiation proteins (Figure 2B-C).

**Figure 2:**
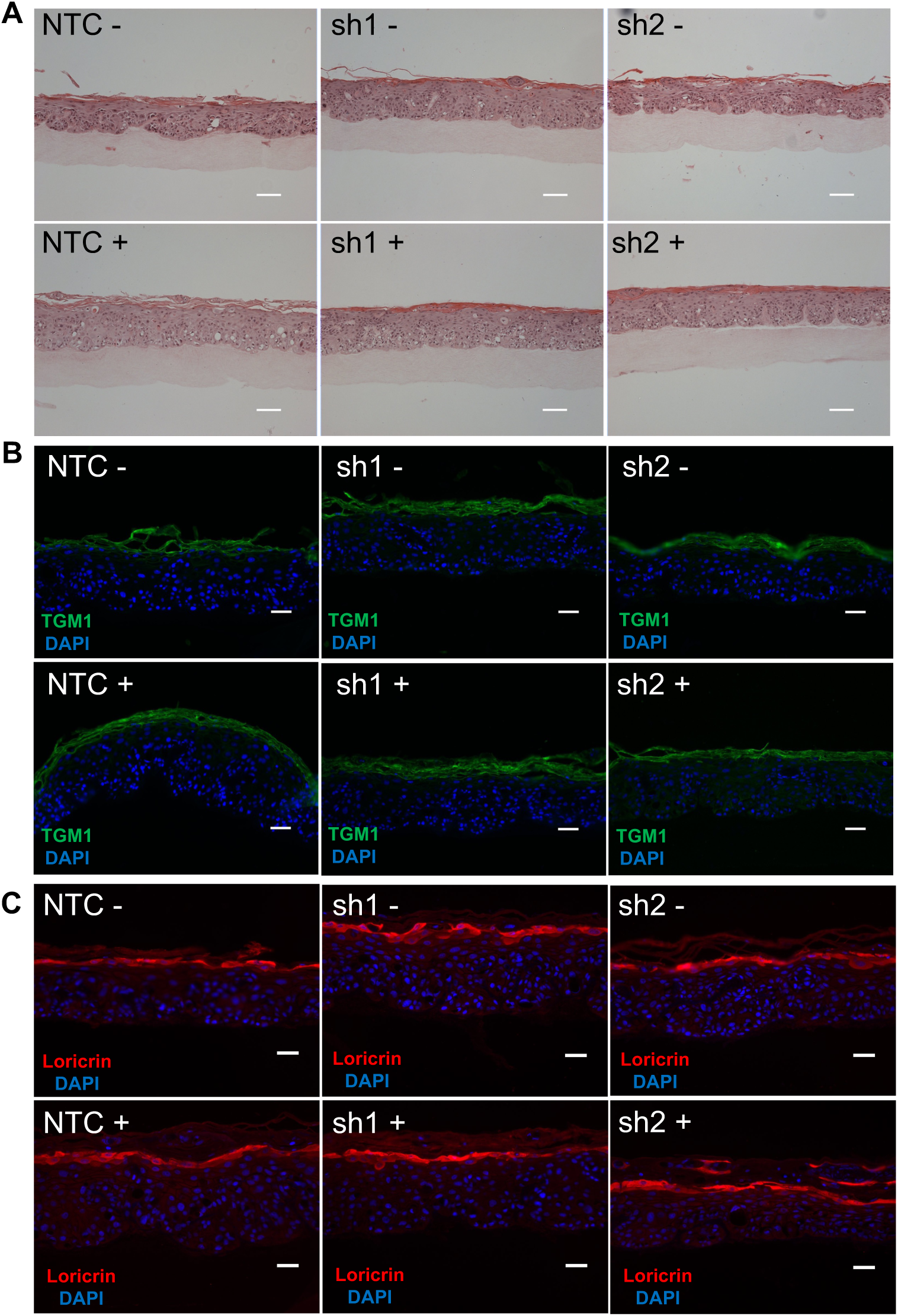
Analysis of stratification and terminal differentiation in 3D HSEs. (A) Representative images of H&E staining for HSEs constructed with NTC, sh1, or sh2 keratinocytes without or with Dox treatment (0.1 μg/ml) for 14 days. Scale bar = 100 μm. (B) Immunofluorescence staining for transglutaminase 1 and (C) loricrin. Scale bar = 20 μm.

Nile Red was then used to detect the levels of non-polar and polar lipids in the epidermal layer^32,33^. Staining for both polar and non-polar lipids localised primarily to the stratum corneum in all models (Figure 3A-B). While the level of non-polar lipids was not altered in response to CerS3 depletion, we observed a significant reduction in the levels of polar lipids for the Dox treated sh2 HSEs, and there was a small but non-significant reduction in polar lipids in the Dox treated sh1 HSEs (Figure 3A-B). Together, these results suggest that induced CerS3 depletion within the HSE model disrupts the polar lipid composition of the stratum corneum independently of keratinocyte terminal differentiation.

**Figure 3:**
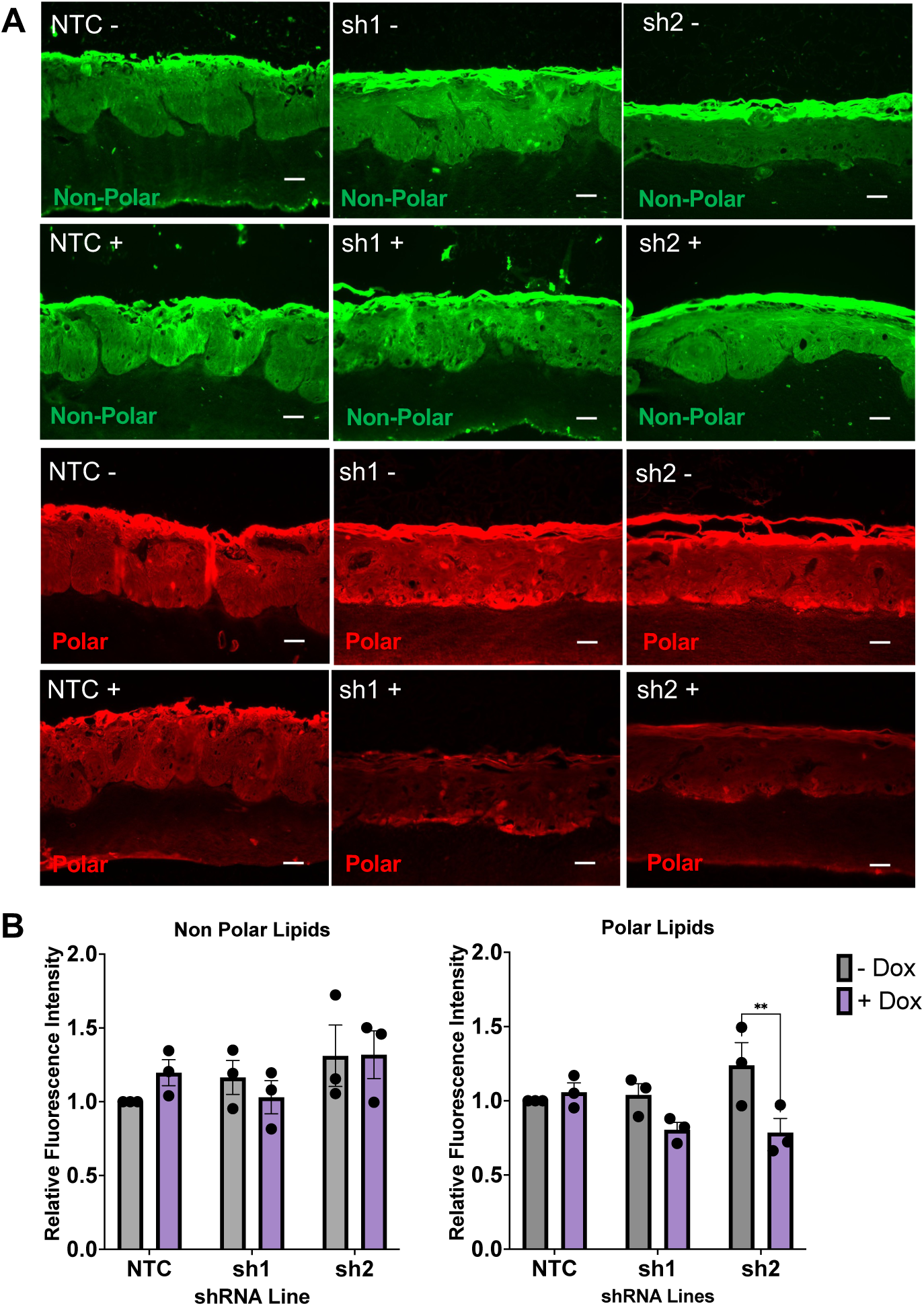
Analysis of lipid content in 3D HSEs. (A) Representative images of non-polar (green) and polar (red) lipid staining using Nile Red, for HSEs constructed with NTC, sh1, or sh2 keratinocytes without or with Dox treatment (0.1 μg/ml) for 14 days. Scale bar = 50 μm. (B) Quantification of Nile Red fluorescence intensity for non-polar (left panel) and polar (right panel) signals for all cell lines without or with Dox treatment. Data represent the mean +/− SEM normalised to the untreated NTC HSEs. **P<0.01 with 2-way ANOVA and Šidák’s multiple comparison test. N=3 experiments.

### CerS3 depletion disrupts the ceramide lipid composition of the HSEs

While Nile Red staining suggested that the lipid composition was perturbed in the CerS3 depleted HSEs, further analysis was required to determine the molecular level changes in the lipidome. We therefore performed liquid chromatography and mass spectrometry (LC/MS) analysis to profile the lipid composition of the HSEs with CerS3 depletion. Overall, LC/MS identified 263 known lipid species across all HSE samples. To assess the similarity between the HSE model and human skin, we compared the proportions of different ceramide classes in the epidermis of our control (NTC) HSEs to the epidermis of normal human skin samples. The percentage of Cer[NH], Cer[NDS], Cer[AH], Cer[AP] and Cer[ADS] in the HSEs were within 5% of those in the donor skin samples. The levels of Cer[NP] were 15% lower in the HSE model compared to human skin, while Cer[AS] and Cer[NS] were 6% and 16.5% higher. respectively. Importantly, Cer[EOS], the most abundant acylCer, was the only acylCer detected in our HSE model at 1% compared to 2% in human skin^36^ (Figure 4A). The overall proportions of ceramides in the epidermis of the human skin samples were also similar to previously reported levels^34,35^. These results therefore confirmed that key epidermal ceramides were present within the HSE model and could be quantitatively analysed by LC/MS.

**Figure 4:**
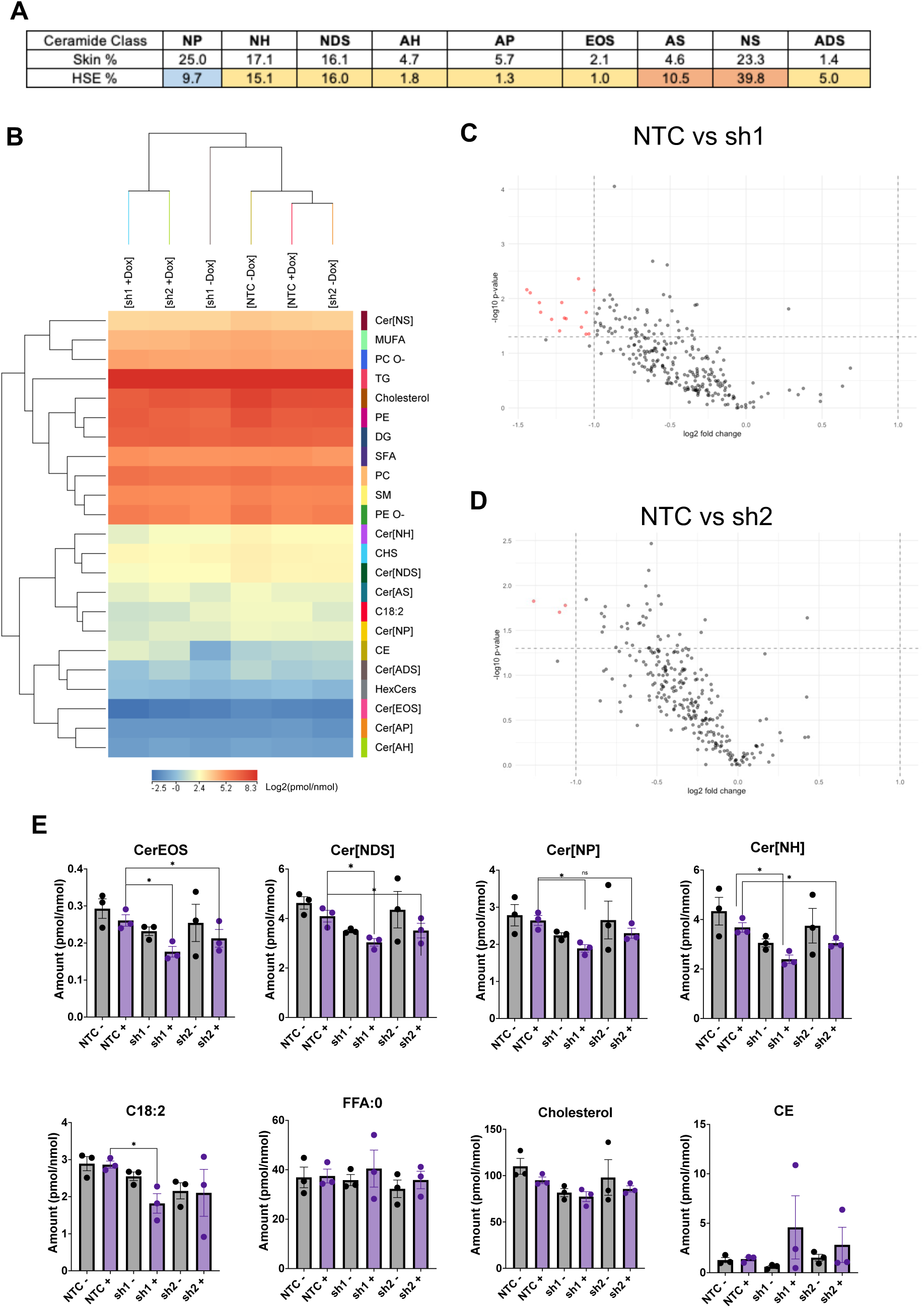
Lipidomic analysis of CerS3 depletion in HSE models. (A) Comparison of the percentage of different ceramide classes out of the total ceramide content between the epidermis of human skin and HSE models constructed with NTC keratinocytes. Yellow = within 5% of human skin, red = higher in the HSE, and blue = lower in the HSE. Data represent the mean of N=3 HSEs from independent experiments. (B) Heat map and hierarchical clustering of lipid classes (263 identified lipid species) and experimental conditions. N=3 experiments. Data represent Log2 transformed levels of each lipid class (pmol) relative to total lipid content (nmol). (C) Volcano plot showing Log2 fold-change and −Log10 P value for all the annotated lipids detected in the HSE models comparing NTC + Dox vs sh1 + Dox and (D) NTC + Dox vs sh2 + Dox. (E) Analysis of ceramide content grouped by different classes, including Cer[EOS], Cer[NDS], Cer[NP], and Cer[NH] alongside levels of linoleic acid (C18:2), saturated fatty acids (FFA:0), total cholesterol, and cholesterol esters (CE), in HSEs constructed with all cell lines without or with Dox treatment for 14 days. Data are represented as mean +/− SEM of pmol of lipid class per nmol of total lipid content. *P<0.05 with unpaired two-tailed Student’s t-test. N=3 experiments.

We next examined the effects of CerS3 KD on lipid composition of the HSEs. Hierarchical clustering based on all identified lipids grouped by lipid class revealed distinct patterns of lipid composition, and the Dox treated sh1 and sh2 samples clustered together and away from the untreated and NTC samples (Figure 4B), indicating a clear effect of CerS3 depletion on the lipid profile of the HSE. CerS3 depletion in both the sh1 and sh2 lines predominantly caused a downregulation of lipids in the HSE model (Figure 4C-D), consistent with the Nile Red staining. Based on a significance criterion of log2 Fc ≥ 1 and a −log10 p value of 1.3, 15 and 3 specific lipid species were significantly downregulated in the Dox treated sh1 and sh2 models, respectively. In addition to individual lipids, we observed a significant downregulation of the ceramide classes Cer[NDS], Cer[NH], and CER[EOS] for both sh1 and sh2 Dox treated HSEs compared to NTC samples, and a downregulation of Cer[NP] for sh1 only (Figure 4E). Linoleic acid (C18:2) is the most common fatty acid used to esterify and create acylCers and is cleaved to form the omega-acylated-hydroxy-Cers (ω-OH-Cer)^7,26^. Linoleic acid was significantly depleted in the sh1 +Dox group (Figure 4E), which we hypothesise may be a feedback response to the decrease in availability of acylceramides. By contrast, there were no significant differences in saturated free fatty acids, cholesterol or other ceramide classes (Figure 4E; Supplementary Figure S3).

Ceramide chain length distribution plays an important role in barrier mechanics^21,25,35,37^, and an increase in shorter chain ceramides leads to a decrease in skin barrier function^37,38^. Therefore, we next analysed ceramide chain length changes within our model. Combining all ceramide classes together, there were significant decreases in longer carbon chain length ceramides (C42 and C44) in both the Dox treated sh1 and sh2 samples and a significant increase in the shorter ceramides (C34) in the sh2 group compared to NTC samples (Figure 5A). Looking at specific ceramide classes, Cer[NS] and Cer[NH] showed significant reductions in the ratio of long to short chain lipids for the sh1 and sh2 compared to NTC (Figure 5B). Together, these findings indicate that CerS3 depletion disrupts the ceramide composition of HSEs, resulting in reduced ceramide content and a shift in the long to short ceramide chain length ratio.

**Figure 5:**
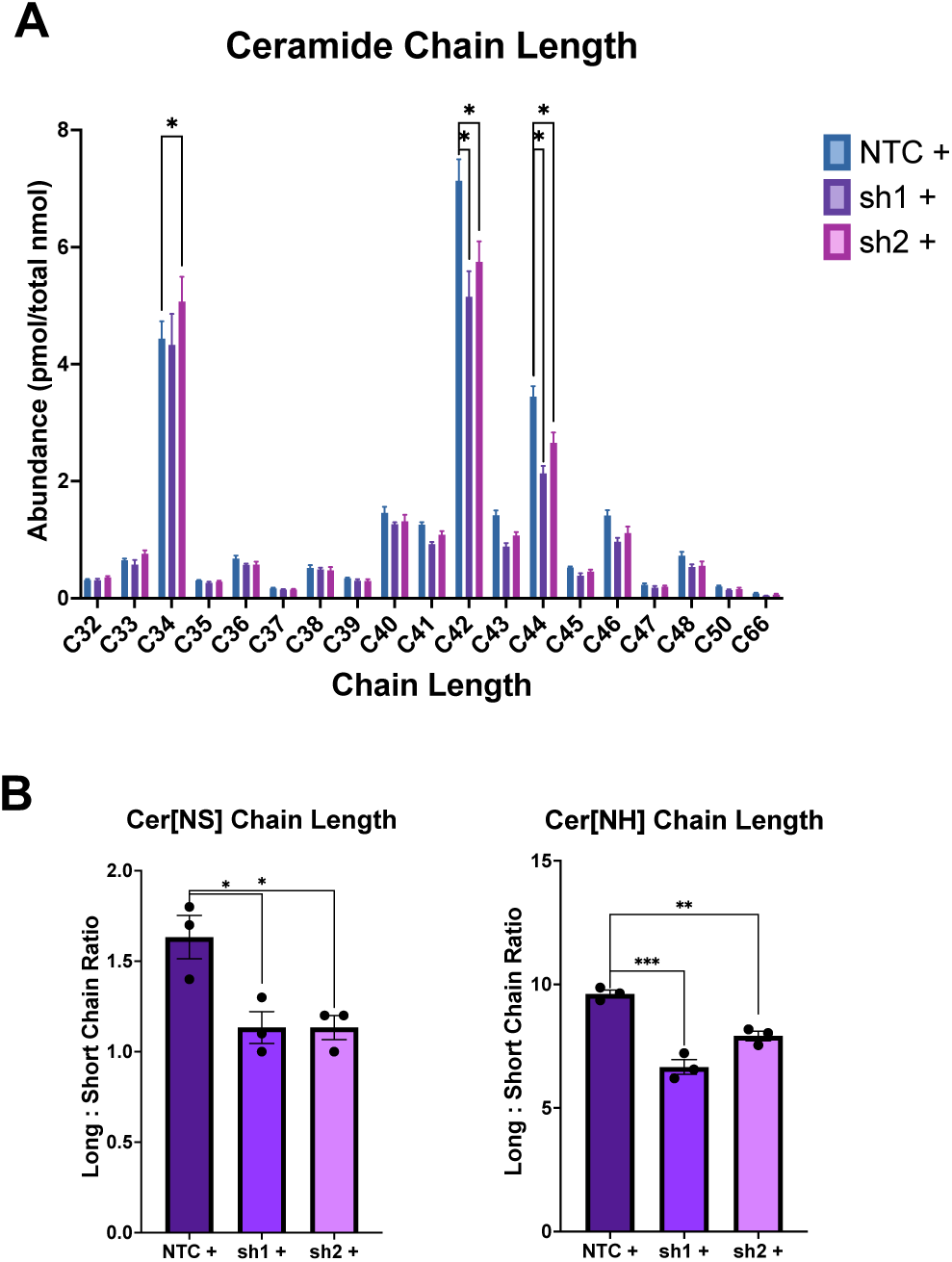
Effects of CerS3 depletion on ceramide chain length in HSE models. (A) Analysis of ceramide chain length in Dox treated HSEs. Data represent the abundance of ceramides with 32 to 66 carbons and are represented as mean +/− SEM. Multiple unpaired t-tests, *P<0.05. N=3 experiments. (B) Molar ratio of long chain ceramides (> 40 carbons) to shorter chain ceramides (<40 carbons) for the most significantly downregulated classes Cer[NS] and Cer[NH]. Data represent the mean +/− SEM. *P<0.05, **P<0.01 and ***P<0.001 with unpaired two-tailed Student’s t-test. N=3 experiments.

### Lipid restoration lags CerS3 protein expression in a model of lipid recovery

A unique feature of the inducible shRNA model is the ability to switch the KD mechanism on and off in a controllable fashion. To further leverage this inducibility we investigated lipid recovery within the HSE model following restoration of CerS3 expression. Time course analysis of CerS3 expression in 2D culture showed that protein levels began to recover within 3 days after Dox removal and were fully recovered within 6 days (Supplementary Figure S4). We then extended this approach to 3D culture, where CerS3 was depleted for 7 days with the addition of Dox and then allowed to recover for 7 days in the absence of Dox. Lipid content in the recovery model was compared to untreated controls or HSEs with CerS3 depletion over the full 14 day culture (Figure 6A). Western blot analysis confirmed a complete recovery of CerS3 at the protein level, in both the sh1 and sh2 HSEs compared to continuous Dox treatment (62% and 55% Cers3 depletion, respectively; Figure 6B-C). As before, the polar lipid content was significantly reduced in the sh2 cultures treated with Dox and slightly, but not significantly, reduced in the sh1 treated cultures. In the sh2 recovery condition, polar lipid content partially recovered to the untreated levels, resulting in non-significant differences compared to the continuous CerS3 depleted condition (Figure 6D-E). These results suggest that there is a latent period between recovery of CerS3 protein expression and full restoration of polar lipid content, and that 7 days is insufficient for a complete recovery of the lipid composition within the in vitro HSE models. Moreover, these findings demonstrate the utility of HSEs with inducible ceramide depletion for the investigation of lipid barrier damage and repair.

**Figure 6:**
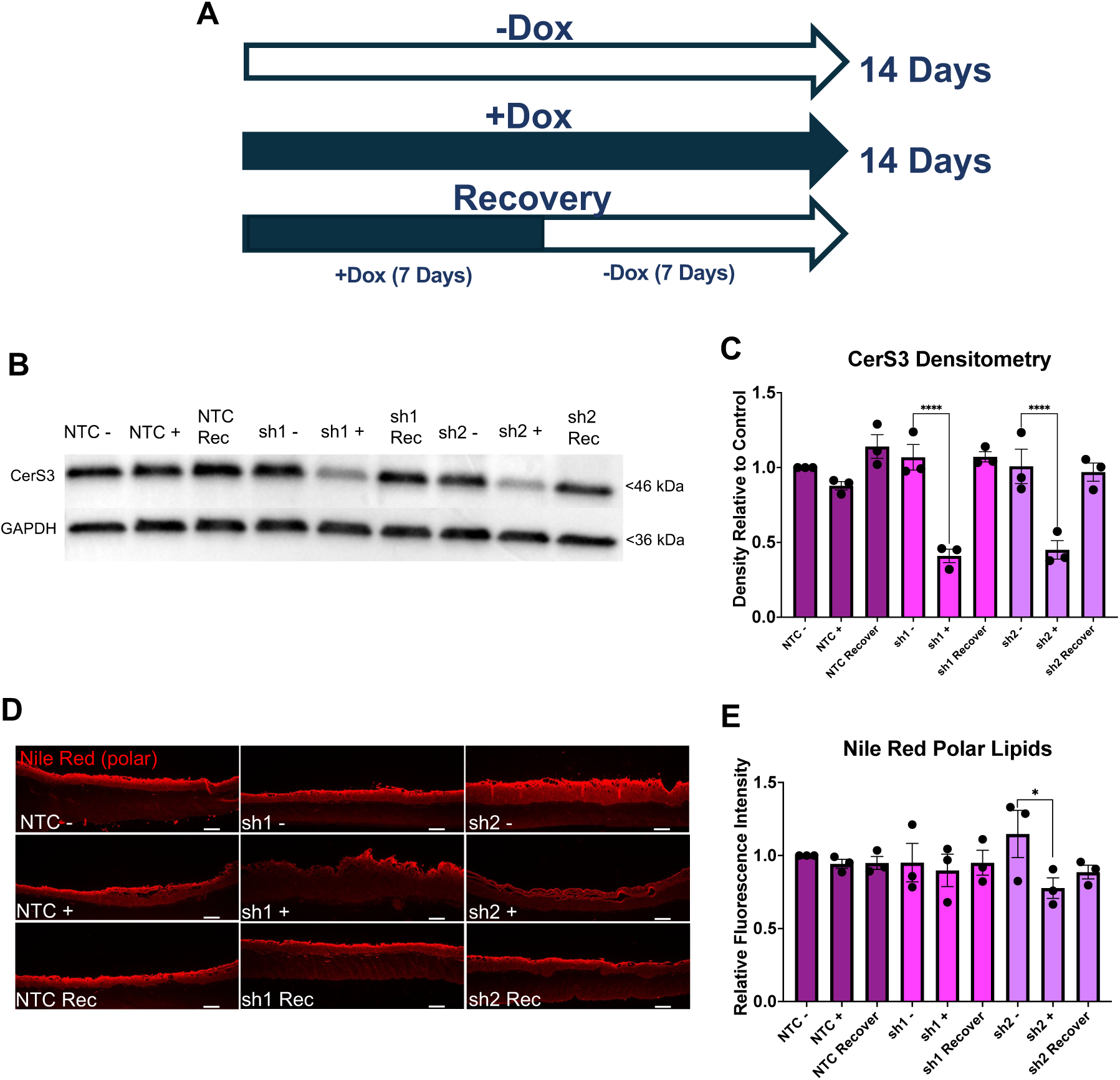
Analysis of CerS3 and lipid restoration in a recovery model. (A) Schematic of 14-day experimental timeline for 3D recovery model, including untreated control (14 days - Dox), sustained knockdown (14 days +Dox), and recovery (7 days +Dox followed by 7 days - Dox). (B) Representative Western blot of CerS3 and GAPDH expression in untreated controls, knockdown, and recovery conditions for NTC, sh1, and sh2 HSEs. (C) Quantification of CerS3 protein levels relative to GAPDH and normalised to NTC untreated controls (-Dox). Data are presented as mean +/− SEM. ****P<0.0001 with 2-way ANOVA and Šidák’s multiple comparison test. N=3 experiments. (D) Representative fluorescence images of polar lipids detected with Nile Red staining in untreated controls, knockdown, and recovery conditions for NTC, sh1, and sh2 HSEs. Scale bar = 100 μm. (E) Quantification of Nile Red fluorescence intensity normalised to NTC untreated controls. Data are represented as mean +/− SEM. *P<0.05 with 2-way ANOVA and Šidák’s multiple comparison test. N=3 experiments.

## Discussion

In this study, we established and characterised a 3D HSE model with tuneable depletion of ceramides through inducible shRNA knockdown of CerS3. We demonstrate that depletion of CerS3 disrupts the lipid composition of the epidermal layer, including a reduction in distinct classes of ceramides and a shift in ceramide chain length ratios, consistent with the known roles of CerS3 in the synthesis of ULC ceramides^7,27,39^. Given that CerS3 levels were not completely knocked down and there were no observable changes in terminal differentiation, we propose that the model best replicates mild forms of lipid deficiency, such as dry skin. By contrast, total loss of CerS3 is known to cause ARCI and more severe phenotypes in skin equivalent models^21^, suggesting that CerS3 levels correlate with the severity of skin disease. Likewise, CerS3 levels are reduced in atopic dermatitis^12,15,25,38^. The ability to tune CerS3 levels in vitro may therefore be a useful approach to gaining new insights into its role in skin barrier diseases.

The inducible shRNA strategy employed here has several notable advantages for modelling lipid disorders in human skin. The efficient induction of transgene expression in over 90% of cells and lack of leakiness provided tight control over CerS3 expression with doxycycline, and as demonstrated in the recovery experiments, could be useful for studying kinetics and mechanisms of epidermal barrier repair. In addition, genetic targeting of individual enzymes, such as CerS3, is advantageous for dissecting the role of specific biosynthetic pathways, and this approach could be applied to other ceramide, fatty acid, or cholesterol synthesis pathways in the future. In this study, we also employed engineered immortalised N/TERT keratinocytes, which were advantageous for lentiviral transduction, scale-up, and consistency, while retaining the ability for form well differentiated HSEs. In future studies, it will be interesting to compare the different effects of CerS3 depletion in HSEs generated with primary keratinocytes, which may display a slightly different lipid profile^40–43^.

Using mass spectrometry, we identified the specific classes of lipids downregulated by CerS3 depletion, namely Cer[NH], Cer[NP], Cer[NDS] and the most abundant acyCer, Cer[EOS]. The former two are notable biomarkers for atopic dermatitis^37^. Our model also recapitulated the frequently reported long to short chain ceramide shift, which compromises barrier function^35,37,38^. While reductions in the abundance of Cer[EOS] were expected based on the known functions of CerS3^8,44,45^, the reduction in the precursor linoleic acid and ceramides in other biosynthetic pathways, suggests that CerS3 depletion has indirect effects on the ceramide composition of the epidermis. The inducible model described here may be useful for dissecting crosstalk and feedback mechanisms of ceramide synthesis in future studies. Likewise, incorporation of functional assays, such as transepidermal water loss, and structural analysis of the LLB with small angle x-ray diffraction^5,13,46^ could provide additional insights into structure-function relationships within the model.

In summary, the inducible HSE model developed here provides a tuneable platform in which to modulate and study lipid synthesis in the skin. We propose that these systems will be useful tools for investigating human-specific mechanisms of epidermal barrier function and disease processes. In addition, they have the potential to support animal-free testing of drugs or skincare products for lipid disorders in human skin.

## Materials and Methods

### 2D Monolayer Culture

All reagents were from ThermoFisher unless otherwise stated. The N/TERT human keratinocyte line, was provided by the Rheinwald Lab^47^. N/TERTs were cultured at 37C and 5% CO_2_ in FAD medium containing a 1:3 mixture of Ham’s F12 and Dulbecco’s modified Eagle’s culture medium, supplemented with 1% (v/v) penicillin/streptomycin (p/s), 1.8×10^-4^ M adenine [Sigma], 10% (v/v) Foetal Bovine Serum (FBS) [Biosera], 5 µg/ml insulin [Sigma], 0.5 µg/ml hydrocortisone, 10 ng/ml epidermal growth factor [Peprotech], 1 x 10^-10^ M cholera toxin [Sigma], seeded in T75 flasks (6,000 cells/cm^2^ seeding density). N/TERTs were passaged at 60-70% confluency by rinsing with pre-warmed Versene solution followed by incubation with pre-warmed 0.25% trypsin-EDTA. Suspension was neutralised with an equal volume of FAD medium before being collected and centrifuged at 200 g for 5 minutes. Cells were counted and replated at a density of 1×10^6^ cell per T75 flask.

Primary human fibroblasts were provided by Prof. Michael Philpott and obtained from healthy donor abdominal skin [ethical reference number LREC 08/H0704/65]. Fibroblasts were seeded at a density of 12,000 cells/cm^2^ and cultured at 37C and 5% CO_2_ in DMEM supplemented with 1% p/s (v/v) and 10% (v/v) FBS. Fibroblasts were used up until passage 7.

### Decellularised Extracellular Matrix Preparation

For the HSE dermal matrix, we used a decellularised extracellular matrix hydrogel (dECM) made from pig skin^31^. Fresh pig skin was cut into 1 cm x 1cm pieces using scalpels with as much fat and hair removed by manual scraping. Pieces were soaked in PBS with 1% (v/v) p/s for 2 hours followed by weighing in sterile tubes. Skin was frozen at −80C for 5 hours prior to being freeze dried overnight. Tissue was digested overnight at 4C in 560 U/L Dispase [Sigma] solution dissolved in a 10 mM sodium acetate (pH 7.5) and 5mM calcium acetate buffer solution at 2 ml per gram of tissue to disrupt the epidermal-dermal association. The epidermis was then scraped off using tweezers and the dermis washed thrice in sterile ddH_2_O, before being washed with constant stirring overnight at room temperature in 70% ethanol. The dermis was then treated with 0.25% Trypsin/0.1% EDTA for 1 hour at 40C using a heated plate stirrer. Dermis was again washed in sterile ddH_2_O, before being incubated at room temperature under constant stirring for 6 hours in a 1% Triton-X, 0.26% EDTA, 0.69% Tris buffer solution. After 6 hours, this solution was replaced with fresh buffer solution and the dermis was incubated overnight on the magnetic stirrer. The tissue was then washed thrice in sterile ddH_2_O, followed by freezing at 80C before being freeze-dried overnight. The dried tissue was weighed and digested at 20 mg/ml acidic pepsin solution, which was prepared by dissolving Pepsin A [Sigma] at a final concentration of 1 mg/ml in 0.01 M hydrochloric acid. The dermis was digested at room temperature for 3 days under constant stirring until tissue was fully dissolved. The resulting dECM was aliquoted and stored at −20°C.

### 3D Human Skin Equivalent Models

HSE models were made in 12-well Transwell plates utilising 0.4 µm inserts [Merck]. Fibroblasts were mixed with thawed aliquots of dECM to make a 1×10^5^ cells/ml suspension and 340 µl of solution was subsequently pipetted onto the apical side of the Transwell insert. The solution was left to solidify for 2 hours in the incubator. After which, 340 µl of FAD- suspended N/TERTs were added on top of the solidified gel. An additional 340 µl of FAD medium was added to the apical side of the Transwell insert, with a further 500 µl of FAD added to the bottom of the well plate prior to overnight incubation. The following day, the inserts were raised to an air-liquid interface by removing medium from the apical side of the Transwell insert, leaving the top surface of the construct dry. HSEs were maintained at an air-liquid interface for 14 days with basal media changes conducted every day. From day 7 onwards, basal media was supplemented with 50 µg/ml ascorbic acid.

### Design and Selection of CERS3 shRNA Sequences

Three lentiviral inducible SMARTvectors (100 µl, 1×10^7^ TU/ml) were purchased for the *CERS3* protein coding targets and the shRNA target sequences were selected from pre-designed sequences [Horizon Discovery Ltd]. For the target gene, three knockdown sequences, named sh1, sh2 and sh3, were all unique in that they targeted multiple splice variants and their alignments with the *CERS3* gene did not overlap. The fourth did not have a targeting nucleotide sequence and was used as a non-targeting control group (NTC). Each construct was transduced into separate N/TERT cell populations, making four transduced cell lines in total (Table 1).

**Table 1:**
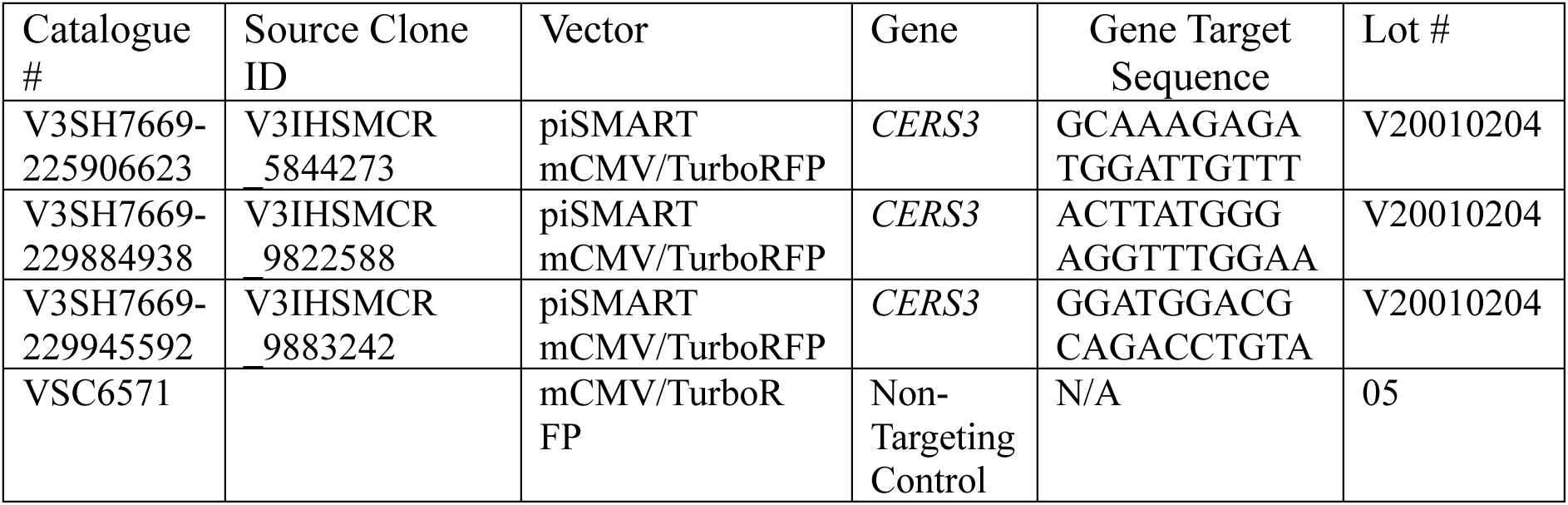
List of the product numbers and identification codes for the selected shRNA constructs.

### Knockdown Cell Line Transduction, Selection and Activation

N/TERT cells to be transduced were seeded at a density of 30,000 cells/cm^2^ in 6 well plates, before incubating in transduction medium with 6 µg/ml polybrene and lentiviral particles at 1.0 multiplicity of infection for 24 hours. Cells were then left to recover in fresh medium for a further 24 hours. After, N/TERTs were cultured until confluent in FAD medium supplemented with 1 μg/ml puromycin before being replated and cultured again until confluent, then cryopreserved. Optimisation of doxycycline concentration was performed on the inducible Non-Targeting Control (NTC) population and monitored by expression of the RFP transgene.

### Flow Cytometry

Transduced cells from monolayer culture were resuspended in PBS (10 x 10^6^ cells/ml) and analysed using the Becton Dickinson FACS Aria IIIu [BD Biosciences]. RFP was detected on the Yellow-Green 582/12nm channel with Forward Scatter (FSC-A) and Side Scatter (SSC-A) detected on the blue laser with doublet discrimination determined by Side Scatter Width (SSC-W).

### Histological Processing, Embedding and Staining

On day 14, HSEs were removed from the Transwell inserts, gently rinsed with PBS and fixed in 4% paraformaldehyde (PFA) for 30 minutes. After two subsequent rinses in PBS, the HSEs were placed individually in tissue processor cassettes and stored in 70% ethanol prior to overnight dehydration in the Tissue-TEK processor [Sakura]. HSEs were paraffin embedded and sectioned using the Leica RM2235 microtome [Leica Biosystems] into 8 μm-thick sections on glass SuperFrost slides.

For H&E staining, sections were incubated in two separate solutions of xylene for 3 minutes each. Afterwards, slides were incubated in separate solutions of decreasing alcohol concentrations for 3 minutes each (100%, 90% and 70%) followed by a further 3-minute incubation in ddH2O. Slides were then sequentially submerged in haematoxylin for 3 minutes, rinsed, then dipped in acid alcohol 5 times before being rinsed in running ddH2O for 5 minutes. Slides were then checked for the presence of stained nuclei under a light microscope, then submerged in eosin solution for 3 minutes before rinsing in running ddH2O for 5 minutes. For dehydration, sections were incubated in increasing solutions of ethanol for 3 minutes each (70%, 90% and 100%), before transferring to the two separate xylene solutions for an additional 3 minutes. Finally, H&E-stained slides were mounted with coverslips using DPX Mountant [Sigma] before imaging with the Nikon Eclipse 80i Stereology Microscope [Nikon].

For immunofluorescence, after deparaffinisation, tissue sections were boiled at 100°C for 15 minutes for antigen retrieval in pH 6 citrate buffer solution consisting of 10 mM sodium citrate tribasic and 1M HCL in ddH2O. Sections were then left to cool in the same buffer for 15 minutes. Tissues were permeabilised with 0.05% Triton-PBS solution for 5 minutes, followed by two subsequent PBS rinses before being placed in a blocking solution of 1% (w/v) fish gelatin [Sigma] and 2% (w/v) FBS in PBS for 1 hour. Sections were then incubated with the selected primary antibodies diluted in blocking solution at 4C overnight. Primary antibodies included anti-keratin 14 (Sc-59724, Santa Cruz [1:100]), anti-Loricrin (ab85679, Abcam [1:80]), anti-TGM1 (HP040171, Atlas Antibodies [1:200]) and anti-tRFP (AB233, Evrogen [1:1000]). The next day, after rinsing with PBS, secondary antibodies diluted in blocking solution were applied to each section, covered then stored for 1 hour at room temperature. Secondary antibodies were anti-mouse Alexafluor 568 (A11004, ThermoFisher [1:1000]) and anti-rabbit Alexafluor 488 (A11008 ThermoFisher [1:1000]). After briefly rinsing with PBS, mounting medium with 4’,6-Diamidino-2-phenylindole dihydrochloride (DAPI) (104139, Abcam) was applied to each of the sections and cover-slipped ready for imaging. Images were acquired using either the Leica DM4000 Epifluorescence or the Leica DM5000B Epifluorescence microscopes [Leica Microsystems]. Normal human skin samples followed the same procedure.

For Nile red staining, samples were first deparaffinised as described previously. Then a stock solution containing 0.05% Nile red in acetone was diluted to 2.5 μg/ml with 75:25 (v/v) glycerol-ddH2O, followed by rapid vortexing. Then a drop of the glycerol-dye solution (20 μl) was applied to each tissue section and immediately covered with a coverslip. Slides were imaged 30 minutes later using the Leica DM4000 Epifluorescence microscope [Leica Microsystems].

### Image Analysis

Fluorescence intensity of IF and Nile red staining were analysed with ImageJ by demarcating the epidermis and measuring the integrated intensity and mean grey values of the epidermis within the region of interest (ROI). For background subtraction, a region without tissue was measured using the same approach. Corrected total cell fluorescence was calculated by multiplying the ROI area by the mean fluorescence of the background and subtracting this value from the integrated density.

### Western Blotting

HSEs were removed from the Transwell inserts and the epidermal layers were separated from the dermal matrix using a scalpel, rinsed with PBS and placed into 1.5 ml microcentrifuge tubes. Tissues were then placed on ice and immersed in 125 µl of a supplemented radio immunoprecipitation assay (RIPA) buffer solution containing 1X protease inhibitor [Sigma]. Samples were vortexed for 20 seconds, homogenised for 20 seconds and sonicated for 5 minutes before being homogenised again for an additional 20 seconds. Samples were then centrifuged for 10 minutes at 4C at a rate of 9,000 xg. Supernatants were collected and stored at −20C.

Equal amounts (20 µg) of protein were loaded onto the 4%−20% Mini-PROTEAN TG Precast Protein running gels [Bio-Rad] and run at 100 V for 1.5 hours. Membrane transfer was performed at 300 A for 1.5 hours onto a 0.45 µm nitrocellulose membrane or the TransBlot Turbo Fast Transfer system [Bio-Rad] was used for the transfer step. Membranes were blocked with 5% milk in Tris-Buffered Saline with 0.1% Tween (TBS-T). Membranes were incubated with primary antibodies, anti-CerS3 [HPA006092 Sigma (1:1000)] and anti-GAPDH [MAB374, Millipore (1:5000)] diluted in blocking solution, overnight at 4C. Membranes were then washed three times in TBS-T for 5 minutes and incubated for an additional 1 hour at room temperature with secondary antibodies, anti-mouse HRP [P0447 Dako (1:15000)] and anti-rabbit [P0448 Dako (1:5000)], diluted in 5% milk. Blots were washed again, three times for 5 minutes, and developed with Enhanced Chemiluminescence HRP substrate. Imaging was carried out on the ChemiDoc XRS imaging system [Bio-Rad]. Densitometry analysis of band intensity was performed using ImageJ.

### RT-qPCR

RNA extraction was performed using the RNeasy Plus Mini Kit [Qiagen], following the protocol provided without the incorporation of either β-mercaptoethanol or dithiothreitol to the lysis RLT buffer. Samples were analysed for RNA content and purity using the NanoDrop ND-1000 Spectrophotometer. For cDNA conversion, RNA samples were diluted to 1 µg total RNA for each sample and the LunaScript RT SuperMix Kit [New England Bio Labs] was used for reverse transcription.

Primers were designed using the University of California Santa Cruz (UCSC) Genome Browser [https://genome.ucsc.edu/] in parallel with the National Centre for Biotechnology Information’s (NCBI) Primer-BLAST tool [https://www.ncbi.nlm.nih.gov/tools/primer-blast/]. The CERS3 primer sequence was (CCATCCAGTAGCTTCGCCTC and TCAGAGAGCAGCTTCCAACG). Primers were subsequently synthesised by Sigma-Aldrich at 100 µM concentrations in solution. GAPDH primers were synthesised by Eurofins [Eurofins Scientific]. PCR reaction master mixes were made on ice using the 2x qPCRBIO Sybr Green Mix Hi-ROX kit [PCR Biosystems], with primers diluted to a 10 µM concentration. For each primer pair, non-cDNA triplicates were used as controls and GAPDH was the housekeeper gene for all samples. Well plates were briefly spun prior to analysis on the StepOne Plu Real-Time PCR System [Applied Biosystems]. Analysis was performed using the ΔΔCt relative quantification method.

### Mass Spectrometry

Lipid extraction was performed using a modified version of the Folch Extraction Method^48,49^. Briefly, HSEs were washed twice in cold PBS before removal of the epidermal layers manually with a scalpel. Epidermal sheets were then placed in Agilent A-Line Amber Glass Vials and dissolved in a 2:1 chloroform/methanol solution at 4C overnight on a shaker. The supernatant was then removed, and the solution was filtered (0.4 µm pore size) and placed into fresh glass vials. Solutions were then dried with nitrogen gas and stored at −80C until analysis.

Lipid extracts were suspended in 200 µL of CHCl_3_:MeOH (2:1 v/v). 40 µL of a mixture of deuterated internal standards was added to the samples to control the intraday heterogeneity and to quantitate the relative abundance of detected lipids. The mixture included deuterated cholesterol-2,2,3,4,4,6-d6 (d6-CH, 320 pmol), deuterated cholesterol sulfate sodium salt (d7-CHS, 160 pmol), Hexadecanoic-9,9,10,10,11,11,12,12,13,13,14,14,15,15,16,16,16-d17 acid (d17-PA, 640 pmol), N-palmitoyl-d31-D-erythro-sphingosine (d31-Cer16:0, 8 pmol), glyceryl Trihexadecanoate-d98 (d98TG 48:0, 160 pmol), 1-pentadecanoyl-2-oleyol(d7)-sn-glycerol (15:0-18:1(d7) DG, 272 pmol), 25,26,26,26,27,27,27-heptadeuteriocholest-5-en-3ß-ol (9Z-octadecenoate) (18:1(d7) CE, 243 pmol), 1-pentadecanoyl-2-oleoyl(d7)-sn-glycero-3-phosphocholine (15:0-18:1(d7) PC, 212 pmol), 1-pentadecanoyl-2-oleoyl(d7)-sn-glycero-3-phosphoethanolamine (15:0-18:1(d7) PE, 225 pmol), N-oleoyl-D-erythro-sphingosylphosphorylcholine-d9 (d18:1-18:1(d9) SM, 217 pmol). The volume was filtered through the Captiva Vacuum (Agilent Technologies) with a pore size of 0.2 µm, and the filtrate was collected in graduated glass vials, dried under nitrogen flow and suspended in 200 µL CHCl_3_:MeOH (2:1 v/v).

For quantitative measurement of cholesterol a GC-MS analysis was performed. 20 µL of the lipid extract was dried under nitrogen and then derivatised with 20 µL of 1% trimethylchlorosilane in pyridine. The reaction carried out at 60C for 60 minutes can produce the trimethylsilyl (TMS) derivatives of most lipids. The GC-MS instrument was an 8890 GC System combined with a 5977B Series MSD single-quadrupole (Agilent Technologies). Helium was used as the carrier gas; the flow rate was 1.2 ml/min. The analysis was conducted on the 30m x 0.250mm (i.d.) x 0.25µm film thickness HP-5MS UI fused silica column (Agilent Technologies), chemically bound with a 5%-phenyl-methylpolysiloxane phase. The GC oven program was as follows: 80C at 0 min, 280C at 33 min, 310C to final run time of 49 minutes. Samples were acquired in scan mode following EI ionization^50^.

Triglycerides, diglycerides, and cholesteryl esters were detected in Reverse Phase-HPLC (RP-HPLC) under positive electrospray ionization ((+)ESI) conditions. Free fatty acids, ceramides, hexosylceramides, and cholesterol sulfate were analysed under negative electrospray ionization ((-)ESI) conditions. To separate polar and hydrophilic compounds, Hydrophilic Interaction Liquid Chromatography (HILIC) was used. Phosphatidylcholines, ether-linked phosphatidyl-choline, sphingomyelins, phosphatidylethanolamines, and ether-linked phosphatidyl-ethanolamine, were detected in HILIC (+)ESI mode. RPLC separation was conducted on an Agilent Technologies Infinity II 1260 series HPLC system equipped with a degasser, a quaternary pump, an autosampler and a column compartment. For the RP-HPLC separation a Zorbax Eclipse Plus C18 column (2.1 x 50 mm, 1.8 µm particle size) (Agilent Technologies) was used. The column temperature was set at 60C, the maximum operating pressure was 600 Bar/9000 psi. Cell extracts were eluted with a binary gradient of (A) MilliQ water (18.2 Ω) and (B) in MeOH/iPrOH 80/20 and the flow rate was 0.3 ml/min. Mobile phase additive was used to improve ionization efficiency. HILIC separation was performed on a HALO HILIC column (Advanced Materials Technolog), 2.1 x 50 mm, 2.7 µm particle size, with maximum operating pressure at 600 Bar/9000 psi. The column temperature was set at 40C. The mobile phase consisted of (A) aqueous solutions of 5mM ammonium formate in MilliQ water (18.2 Ω) and (C) ACN. The mobile phase flow rate was 0.4 ml/min.

The HPLC instrument was connected by an ESI Dual Agilent Jet Stream (AJS) interface with an Agilent Technologies (Santa Clara, CA, USA) 6545 Quadrupole Time of Flight (Q-TOF) as mass spectrometer. Nitrogen was used as a gas for both the nebulisation and desolvation processes. The ion source gas temperature was 200 °C with a flow rate of 12 L/min; the nebuliser pressure was 40 psi. Sheath gas temperature was set at 350 °C; sheath gas flow rate of 12 L/min. The fragmentor voltage parameter was 120 V and the skimmer 40 V. Data were collected in all ion MS/MS modes, at 0, 20, 40V of collision energies. The m/z range for MS an MS/MS of 59-1700 at a mass resolving power of 40,000. Internal mass calibration for accurate mass measurement used two reference ions: m/z 121.0509 and m/z 922.0098 (+)ESI, m/z 119.0363 and m/z 940.0015 (-)ESI.

For analysis, all data were derived by normalizing the response of the individual lipid by the response of the same-class labelled internal standard (e.g. free fatty acids vs deuterated-palmitic acid, cholesterol vs deuterated-cholesterol, etc.) and multiplied by the iSTD pmole. Resulting pmoles were normalized by total nmole amount of each sample. A t-test was performed to determine significant differences between sample groups and how they are related. Differences were considered statistically significant with P-values (p) ≤ 0.05. Additional analyses and graphing (volcano and PCA plots) were conducted using R Statistical Software (v2023.06.1+524 “Mountain Hydrangea”).

## Data Availability Statement

All data will be made available on request.

## Conflict of Interest Statement

The authors state no conflict of interest.

## Acknowledgements

This work was funded by a PhD studentship from the BBSRC London Interdisciplinary Doctoral Programme (LIDo). We thank Dr Matthew Caley and Prof. Edel O’Toole for their advice and guidance on retroviral transductions and HSE methods.

## Author Contributions

Investigation: DD, MM, HK. Formal Analysis: DD, MM, HK, CE, JTC. Writing – original draft: DD and JTC. Writing – review and editing: DD, CE, JTC. Conceptualisation: JTC. Supervision: JTC. Funding Acquisition: JTC

---

**Supplementary Figure S1:**
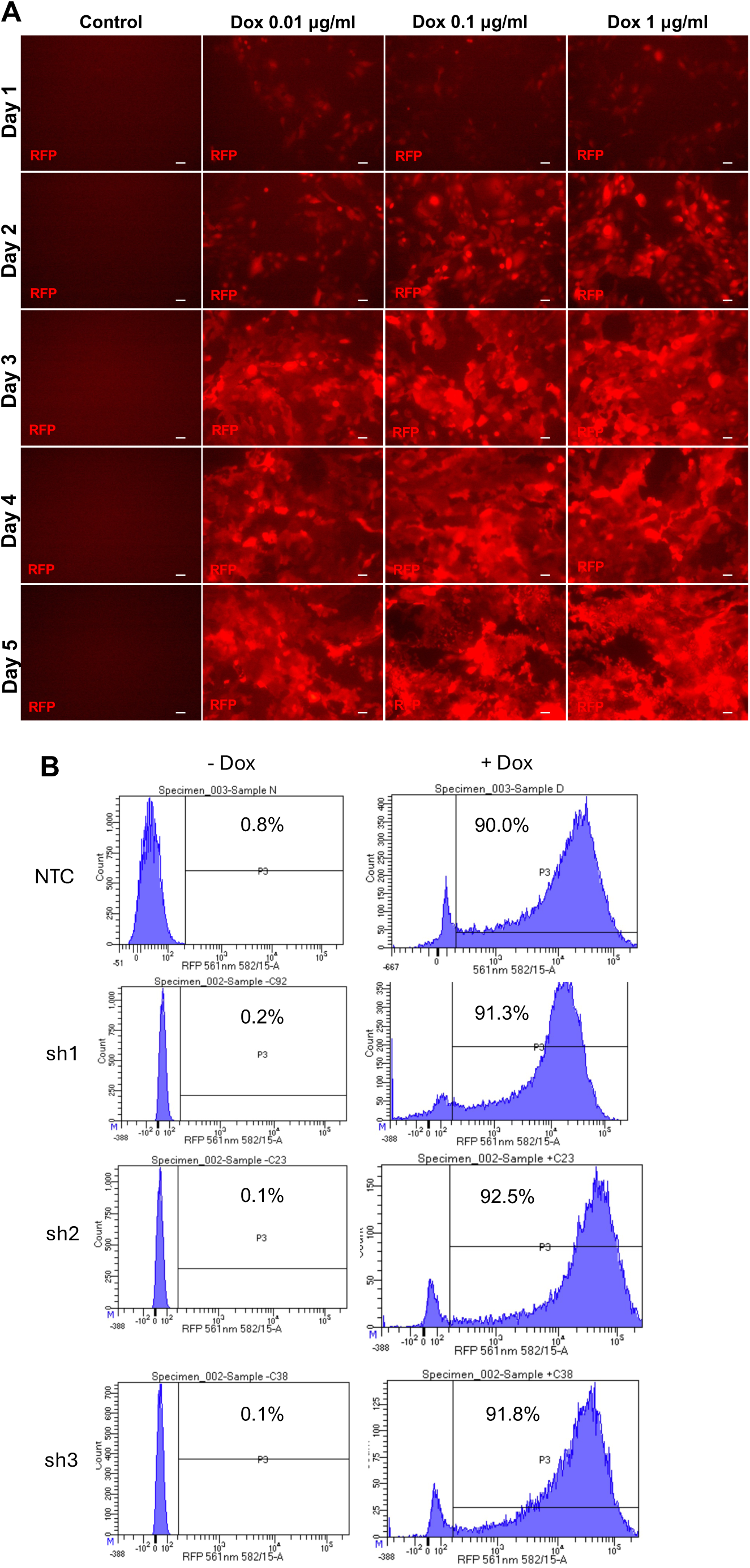
Optimisation of transgene induction with doxycycline. (A) RFP fluorescence imaging in N/TERT keratinocytes transduced with NTC shRNA and treated with 0, 0.01, 0.1 or 1 μg/ml of doxycycline for a period of 6 days. Images are representative of N=3 experiments. Scale bar = 50 μm. (B) Flow cytometry for RFP reporter expression across non-targeting control (NTC) and shRNA knockdown cell lines (sh1, sh2 and sh3) in 2D culture without or with 0.1 μg/ml doxycycline treatment for 7 days. Percentage of RFP positive cells are noted for each condition. Data represent N=1 experiment.

**Supplementary Figure S2:**
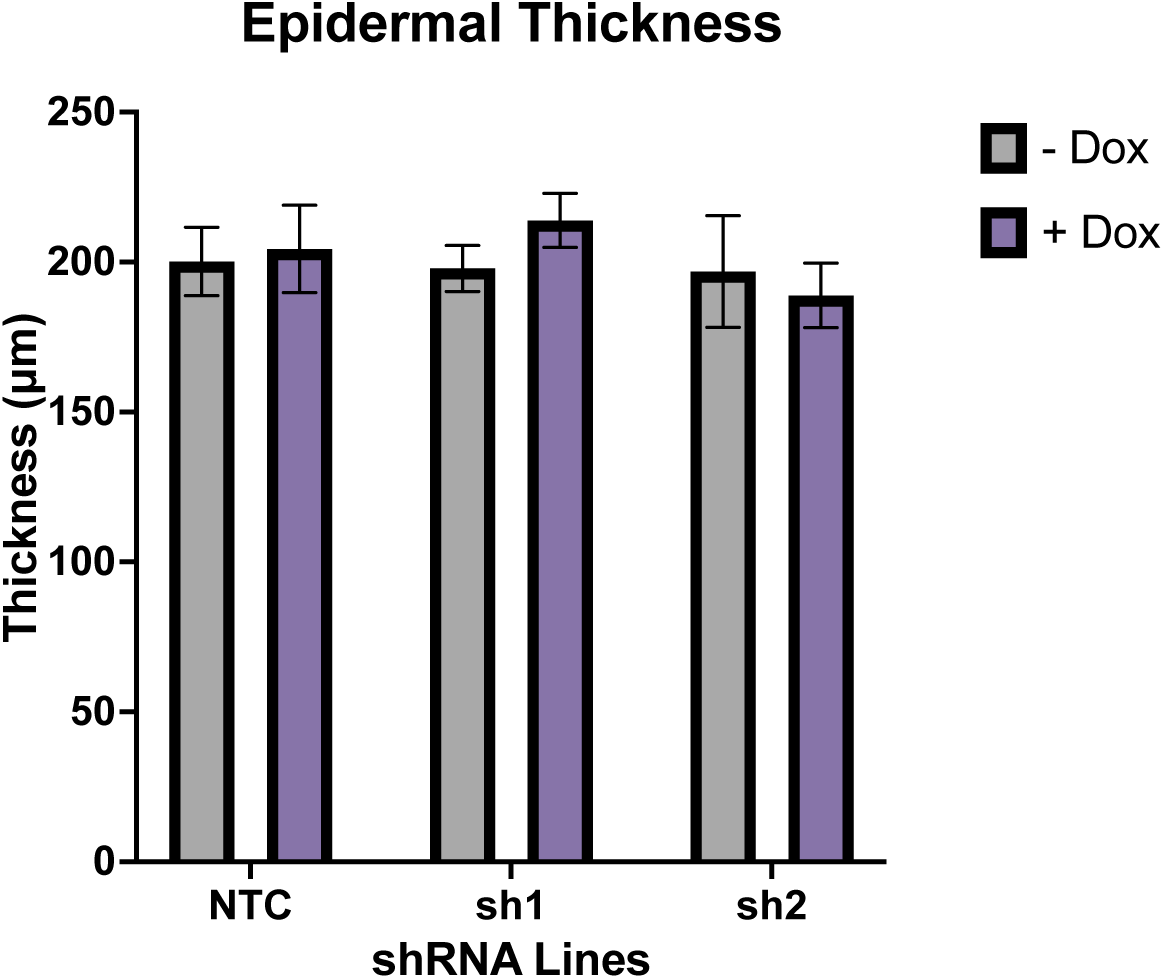
Epidermal thickness. Quantification of epidermal thicknesses using H&E images from HSEs constructed with NTC, sh1, and sh2 lines and without or with doxycycline treatment for 14 days. Data represent the mean +/− SEM of 3 technical replicates and N = 3 independent experiments.

**Supplementary Figure S3:**
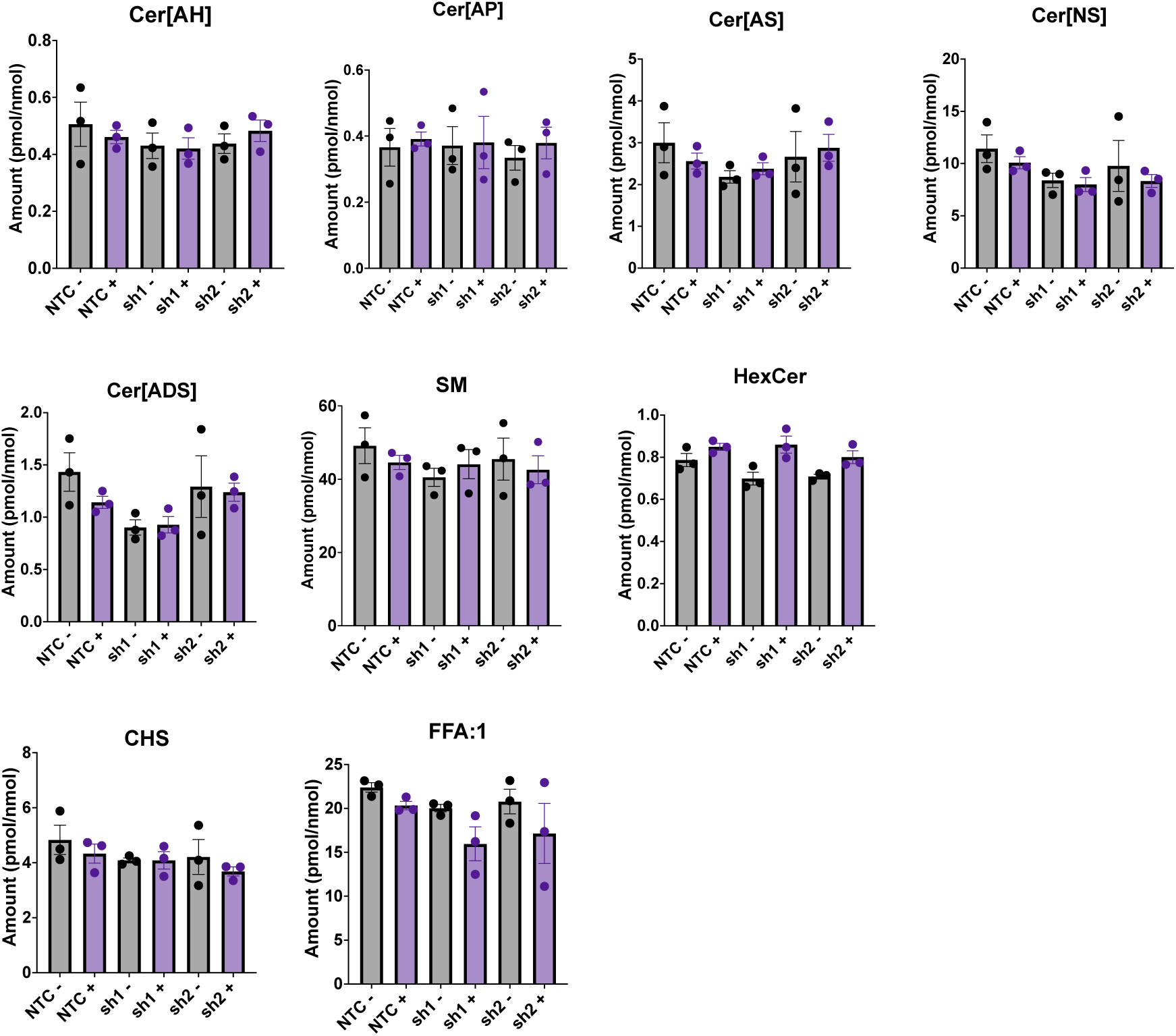
Analysis of additional lipid classes. Quantification of levels of lipids in Cer[AH], Cer[AP], Cer [AS], Cer[NS], Cer[ADS], sphingomyelin (SM), Hexosylceramide (HexCer), cholesterol sulphate (CHS), and unsaturated free fatty acids (FFA:1). Data are represented as mean +/− SEM of pmol of each lipid class per nmol of total lipid content. *P< 0.05 with unpaired two-tailed Student’s t-test. N = 3 experiments.

**Supplementary Figure S4:**
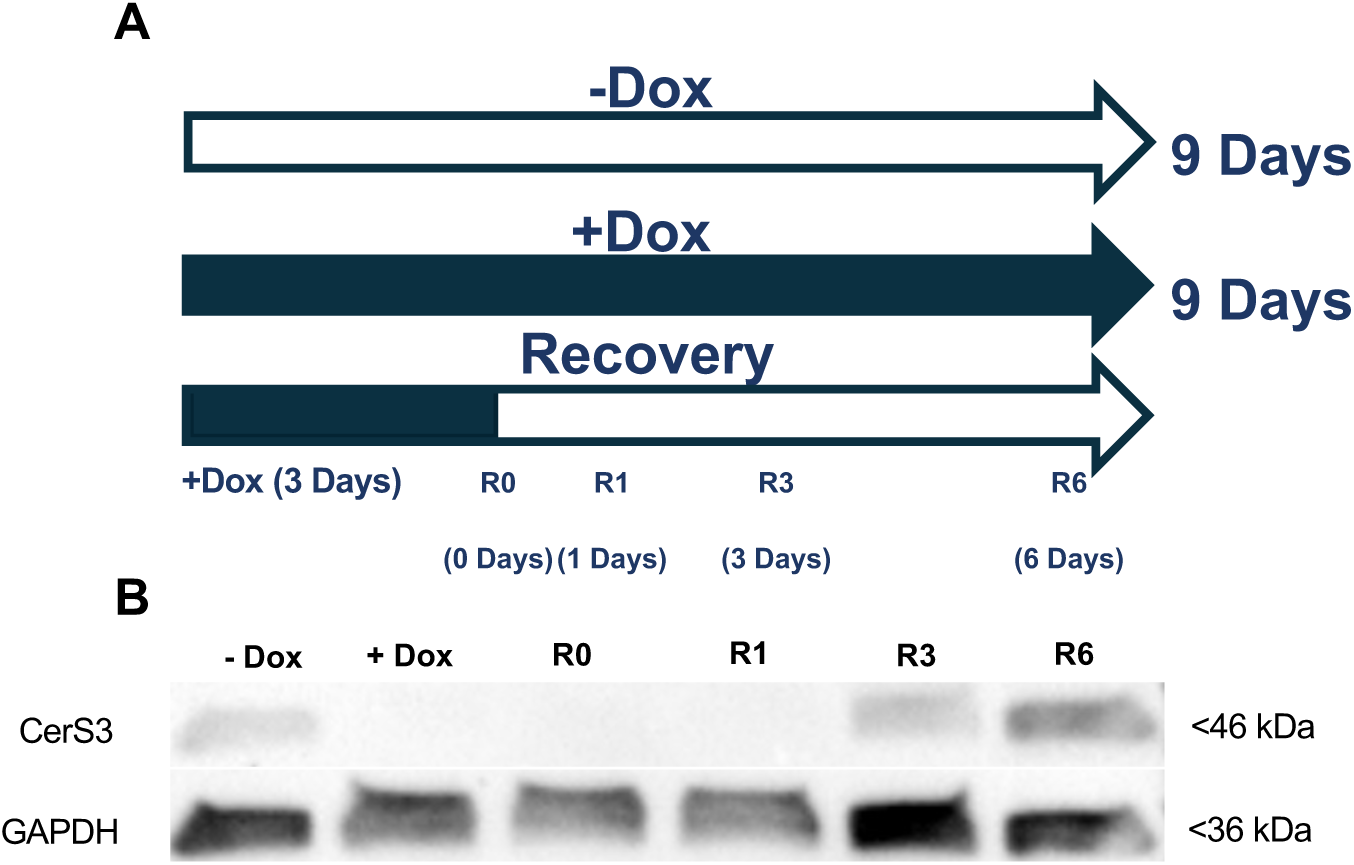
Analysis of CerS3 recovery kinetics in 2D culture. (A) The 2D knockdown recovery experiment culture timeline. The experiment consisted of a -Dox and a+Dox condition for 9 days, plus four independent recovery conditions where cells were treated with Dox for 3 days following by sample collection at 0, 1, 3, and 6 days after removal of Dox (R0, R1, R3 and R6). (B) Representative Western blot for CerS3 and GAPDH for the two controls and four recovery time points.

## Notes

### Competing Interest Statement

The authors have declared no competing interest.

